# Telomere length is highly heritable and independent of growth rate manipulated by temperature in field crickets

**DOI:** 10.1101/2020.05.29.123216

**Authors:** Jelle Boonekamp, Rolando Rodríguez-Muñoz, Paul Hopwood, Erica Zuidersma, Ellis Mulder, Alastair Wilson, Simon Verhulst, Tom Tregenza

## Abstract

Many organisms are capable of growing faster than they do. Restrained growth rate has functionally been explained by negative effects on lifespan of accelerated growth. However, the underlying mechanisms remain elusive. Telomere attrition has been proposed as a causal agent and has been studied in endothermic vertebrates. We established that telomeres exist as chromosomal-ends in a model insect, the field cricket, using terminal restriction fragment and *Bal 31* methods. Telomeres comprised TTAGGn repeats of 38kb on average, more than four times longer than the telomeres of human infants. *Bal 31* assays confirmed that telomeric repeats were located at the chromosome-ends. We tested whether rapid growth is achieved at the expense of telomere length by comparing crickets reared at 23°C with their siblings reared at 28°C, which grew three times faster. Surprisingly, neither temperature treatment nor age affected average telomere length. Concomitantly, the broad sense heritability of telomere length was remarkably high at ~100%. Despite high heritability, the evolvability (a mean standardized measure of genetic variance) was low relative to that of body mass. We discuss the different interpretations of these scaling methods in the context of telomere evolution. It is clear that some important features of vertebrate telomere biology are evident in an insect species dating back to the Triassic, but also that there are some striking differences. The apparent lack of an effect of growth rate and the total number of cell divisions on telomere length is puzzling, suggesting that telomere length could be actively maintained during the growth phase. Whether such maintenance of telomere length is adaptive remains elusive and requires further study investigating the links with fitness in the wild.

## 1 INTRODUCTION

Genetic factors and early life developmental conditions affect growth, with long-term consequences for adult health and fitness, but the underlying proximate mechanisms remain elusive (Dmitriew 2011). Biomarker variables could shed light on causal mechanisms, in particular when manipulated growth rate is reflected in the biomarker and when the biomarker variable in turn is predictive of fitness (Frangakis & Rubin 2002; Boonekamp *et al.* 2017). Telomere length is advocated as a useful biomarker in this context, e.g. Monaghan & Ozanne (2018).

Telomeres are highly conserved nucleoprotein structures that form protective caps at the chromosome-ends where they prevent catastrophic end-to-end chromosome fusion (Armanios & Blackburn 2012). Vertebrate telomeres comprise tandem repeats of a 5’-TTAGGG-3’sequence protected by shelterin proteins (Blackburn 1991). At the chromosome level, telomeres shorten with each cell division through incomplete DNA replication (Olovnikov 1973), single strand breaks induced by reactive oxygen species molecules (Zglinicki 2002), and loss of the single strand overhang (Stewart *et al.* 2003). Telomeres may be restored through the action of telomerase, but the activity of this enzyme rapidly diminishes during early development and it is thought to be mainly inactive in somatic tissues (Hug & Lingner 2006). Consequently, telomeres shorten with age, with the fastest telomere erosion observed during growth (Zeichner *et al.* 1999; Salomons *et al.* 2009). In adult humans, telomere shortening is related to stress (Epel *et al.* 2004; Entringer *et al.* 2011), disease (Calado & Young 2009), and lifestyle factors (Lin *et al.* 2012) and telomere length predicts mortality risk (Boonekamp *et al.* 2013). Telomere length also predicts mortality (Wilbourn *et al.* 2018) and fitness (Eastwood *et al.* 2019) in natural populations of vertebrates and the rate of telomere attrition is correlated to lifespan across species (Tricola *et al.* 2018). Gradual telomere erosion with growth and/or age could reflect a multilevel process of cell division, damage, and repair, linked to fitness. It is therefore intuitive to consider telomere length at the end of the growth period and/or the rate of telomere attrition during growth as candidate biomarkers linking growth to fitness, a supposition supported by earlier studies, e.g. Heidinger *et al.* (2012); Boonekamp *et al.* (2014).

The proximate mechanism of telomere length as a biomarker of the cost of growth is based on the premise that growth acceleration increases the number of telomere base pairs that are lost per cell division (Monaghan & Ozanne 2018). Dependence of the rate of telomere attrition on growth rate could for example arise by increased oxidative stress associated with growth acceleration (Alonso-Alvarez *et al.* 2007; Smith *et al.* 2016), but see Chatelain *et al.* (2019). However, because both the total number of cell divisions and the rate at which cells divide may contribute to telomere loss, experiments are required that disentangle the effects of size and growth rate on telomere length. Only a few studies have attempted to tackle this and they show mixed results: In common terns, incubator temperature affected the time until hatching without affecting size at hatching, and faster growth corresponded with reduced telomere length of hatchlings (Vedder *et al.* 2018). In contrast, a similar experiment in Atlantic salmon larvae found no association between growth rate and telomere length (McLennan *et al.* 2018). We are unaware of other experimental studies attempting to distinguish the effects of size and growth rate on telomere length. Hence whether growth acceleration reduces telomere length remains unclear.

The relationship between telomere dynamics and fitness has been studied almost exclusively in vertebrates, but see Jemielity *et al.* (2007). However, telomerase-based maintenance of the integrity of chromosome-ends appears to be almost ubiquitous in metazoans, with the exception of dipterans (Gomes *et al.* 2010). In insects with terminal telomeres the predominant telomeric sequence is 5’-TTAGG-3’, which is only one guanine nucleotide short of the vertebrate sequence, and many insects appear to express telomerase during development (Korandová *et al.* 2014). Furthermore, fluorescence in situ hybridization (FISH) reveals that insect TTAGG repeats occur almost exclusively at the chromosome ends (Okazaki *et al.* 1993; Frydrychová *et al.* 2004; Vítková *et al.* 2005), suggesting that insects are virtually free from interstitial telomeric repeats. These findings raise the question of whether telomere length is a biomarker of growth and fitness in insects. If confirmed, telomere length could be used as study endpoint, eliminating the need to longitudinally monitor insects which are notoriously difficult to follow in the wild.

As ectotherms, insects have enormous developmental plasticity and as such are powerful systems for studying the effects of developmental conditions on growth and fitness (Richard *et al.* 2019). Growth rate can be manipulated through modification of temperature in ectotherms (Gilbert & Raworth 1996), and because environmental temperatures are variable it can be assumed that insects show an evolved response to temperature variation. Temperature is expected to have differing effects in ectotherms versus endotherms where internal body temperature is maintained.

Endotherms have evolved to operate at a narrow range of temperatures, so attempts to manipulate these are likely to cause acute stress. Ectotherms have evolved to tolerate varying temperatures, and are not expected to attempt to compensate, for instance by increasing food consumption and metabolic rate, in the same magnitude as endotherms, although costs of a modified growth trajectory have been observed in sticklebacks (Lee *et al.* 2010; 2016). However, until a critical upper limit is reached, temperature inevitably increases metabolic rate, increasing both oxygen consumption (Roe *et al.* 1980) and oxidative stress in insects (Lalouette *et al.* 2011; Das *et al.* 2018) and we therefore assume that accelerated growth through temperature will reduce lifespan in insects.

Here, we use the field cricket *Gryllus campestris* to study the effects of temperature and growth on telomere length. Previous FISH studies show that telomeres exist as chromosome-ends in two related cricket species (*G. assimilis* and *Eneoptera surinamensis*) (Palacios-Gimenez *et al.* 2015). We verified that there are telomeric repeats at the chromosome-ends in *G. campestris*, using a combination of approaches (see methods). Subsequently, to study the effects of growth rate on telomere length, we used an experimental design that allows us to disentangle the effects of age and temperature on mass and telomere length. We predict that growth acceleration by temperature results in progressively shorter telomeres with longer exposure to higher temperature.

## 2 MATERIAL AND METHODS

### 2.1 Cricket collection and breeding

We collected adult and subadult crickets from ten sites in Northern Spain (see Table 1) in late April and early May 2018 using burrow traps (for details on trapping see Rodríguez-Muñoz *et al.* (2019a)). Five sites were at low altitude (84 ± 59m above sea level (mean ± SD)) and five sites were at high altitude (1148 ± 164m). Captured crickets were maintained in a research facility in the study area (near location L3, see Table 1), in individual 7×7×10cm (LxWxH) metal mesh cages (pen holders) in a single temperature-controlled room at 25°C with a 7am-10pm light cycle, reflecting the day length in their source locations. The position of the boxes was rearranged at least weekly to ensure that potential temperature gradients within the room could not systematically affect populations. Once individual females had been adult for at least 3 days they were mated to a single male from their own population by placing the pair together in a 9-cm circular plastic container and observing them until mating occurred, typically within an hour (n=58). In a further 7 cases the pair failed to mate within 1 h, and the female was provided with a 2^nd^ male within the next 2-3 days with whom she mated. After mating, females were provided with a 3-cm petri dish with wet sand for egg laying. Dishes were checked and eggs removed every other day. Eggs were transferred to a new petri dish containing cotton wool moistened with water and maintained at 25°C in an incubator (1-3 petri dishes per female). Females typically laid between a few tens to a few hundreds of eggs each, which were spread out evenly on the surface of the cotton wool to reduce the potential for mould to spread. As nymphs hatched over a period of ~ 8 days, they were transferred to 7×7cm boxes and sent by air to the University of Groningen (the Netherlands).

**Table 1.**
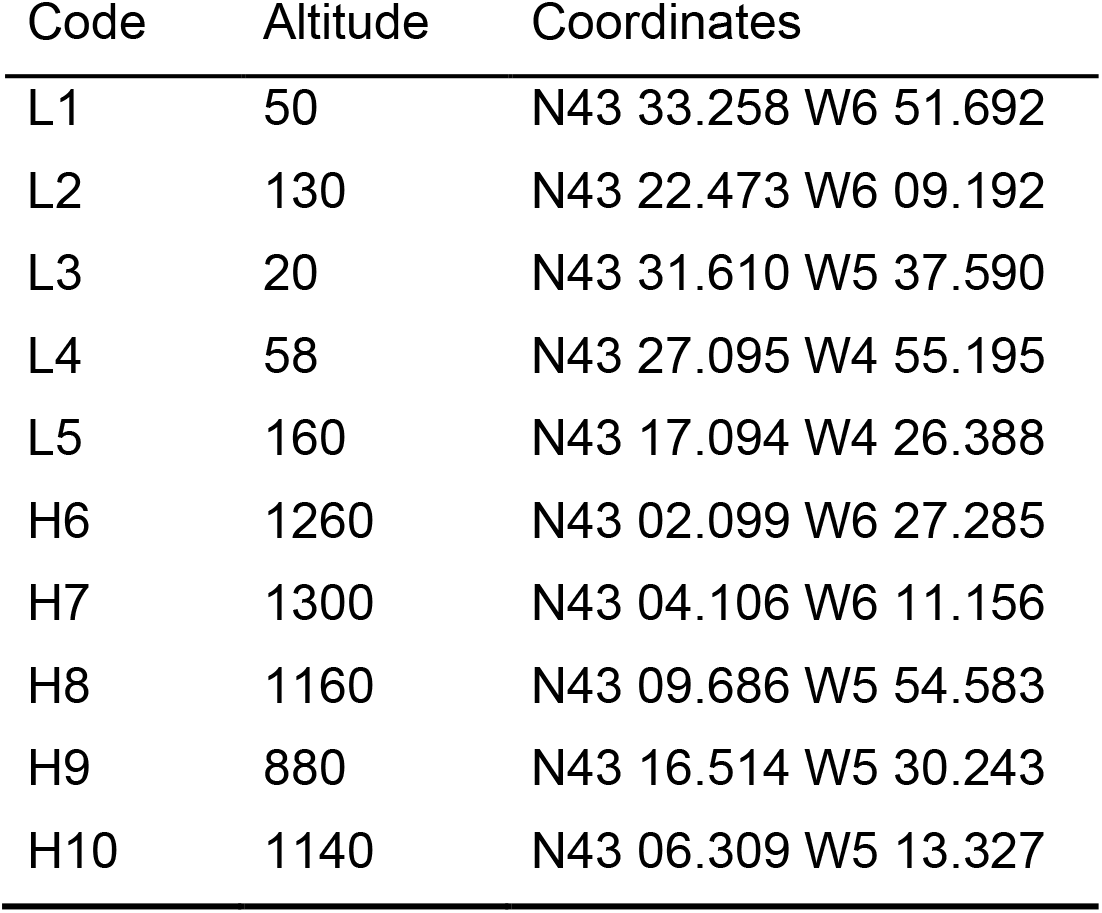
Sampling locations in Northern Spain

### 2.2 Temperature treatment

At the University of Groningen, we housed the nymphs in four identical climate rooms to control temperature, humidity, and the daily light cycle. We set the temperature of two of the four rooms to 28°C and the other two rooms to 23°C, whilst relative humidity was maintained at 50%. All four rooms were on a 12hr light / dark cycle. We took a median of 20 nymphs from each dam (range 6 – 115) and divided siblings over the four climate rooms such that each room had representatives of each family. To minimize potential negative effects of density, up to 10 siblings (median ± SD = 6 ± 4.4) were kept in single boxes of 10 × 10 × 6 cm; we used multiple boxes for large families.

We snap-froze nymphs in liquid nitrogen at 3 different time-points to create “same chronological age” nymphs that differed in their body size due to the temperature treatment, as well as “same size but different chronological age” nymphs. The freezing of the three batches was timed as follows: We sacrificed half of all the boxes of nymphs in both treatment groups after 65 days of treatment. The second half of the 28°C group was sacrificed after 85 days of treatment, and the second half of the 23°C group after 125 days, with the intention that these sample groups had grown to the same size. This design allowed us to disentangle the effects of age and temperature on body mass (see fig 1 for a schematic overview). We measured body mass immediately before snap-freezing and nymphs were stored at −80 °C until DNA extraction for telomere length measurement. In total 545 nymphs from 59 dams survived to sampling, with n=188 (day 65 @ 23°C); n=195 (day 65 @ 28°C); n=53 (day 125 @ 23°C); and n=109 (day 85 @ 28°C). We measured telomere length in a subset of 77 randomly selected nymphs from 29 dams. They were selected at random, per treatment per female when available, but with post-selection verification that the body-mass distributions were not skewed (see Fig. S1).

**Figure 1.**
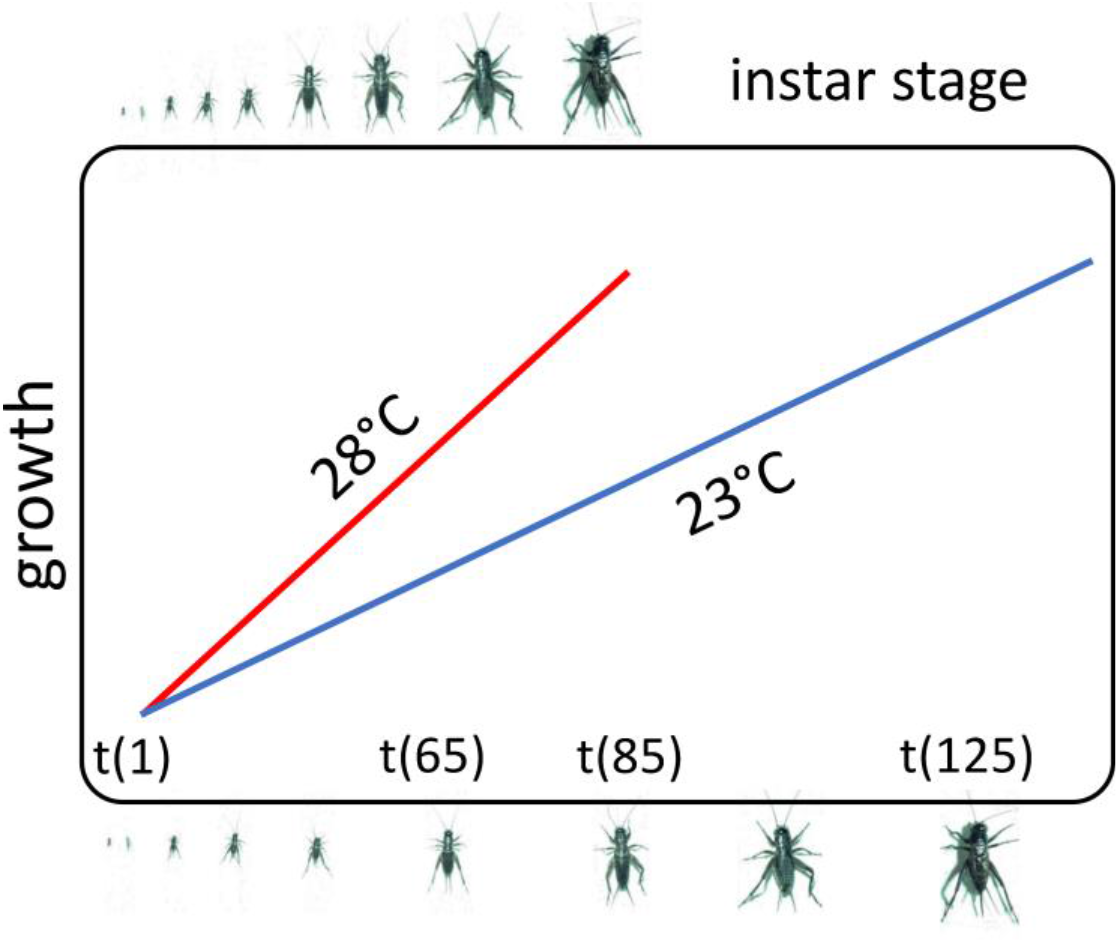
Schematic overview of the experimental design. Early stage nymphs entered temperature treatment conditions at t(1). Temperature conditions (28°C & 23°C) were replicated using a duplicate climate room setup. For each family, siblings were divided over the four climate rooms such that each family was represented in each room by at least one sibling. Half of the nymphs in both temperature conditions were weighed and snap frozen after 65 days of treatment. This group represents same age, different state siblings. The second half of the 23°C group were sacrificed after 125 days of treatment with the aim of achieving the same state relative to 28°C, but different chronological age siblings. Because these two groups of 65 days @ 28°C and the 125 days @ 23°C groups differ in two dimensions, both in time and temperature, we included a third group of 28°C that was sacrificed at an intermediate age (t85), thus achieving greater statistical power to disentangle the effects of temperature from the effects of age.

### 2.3 DNA extraction

DNA was extracted for telomere length measurements. Up to 100mg of thorax-tissue was obtained from frozen specimens and prepared for DNA-extraction using the Blood & Cell Culture DNA Midi Kit (#13343) by Qiagen, using 100/G Genomic-tips, following the manufacturer’s protocol described in Qiagen Genomic DNA Handbook. DNA concentrations were measured in duplo with a Nanodrop 2000c from Thermo. Solutions were scaled to a allow a maximum of 4μg genomic DNA to fit homogeneously into a 100μl agarose plug, using the CHEF Genomic DNA Plug Kit (#170-3591) by Bio-Rad Laboratories. Only ¼ of each agarose plug was used for telomere length analysis.

### 2.4 Nuclease *Bal 31* assay

We carried out a *Bal 31* assay on six individual adult field crickets that were not part of the temperature experiment in order to establish that telomeric repeats occurred at the chromosome-ends. This assay rests on the premise that *Bal 31* degrades DNA from the chromosomal ends and hence if *Bal 31* exposure reduces telomere length then this is demonstrative of their end-cap position (Atema *et al.* 2019). A positive result however does not rule out the coexistence of interstitial telomeric repeats, which can potentially be substantial in number and highly variable among individuals. The outcome of the *Bal 31* assay is therefore particularly informative when the assay is done using denatured DNA: long interstitial telomeric repeats will then be revealed by the telomere probe binding to all TTAGG sequences on the genome, including interstitial repeats (rather than the probe’s affinity being restricted to binding to the single strand telomere overhang of intact double stranded DNA). We therefore carried out the *Bal 31* assay and denatured DNA after gel electrophoresis (see section 2.5 below). *Bal 31* digestion was performed in agarose plugs containing isolated genomic DNA. Prior to the digestion, plugs were incubated for 1 hour with the *Bal 31* 1× reaction buffer (supplied by manufacturer). Each agarose plug was cut into 4 equal pieces in separate reaction tubes. One piece (T=0) was directly washed on ice 3× for 10 minutes with 50mM EDTA followed by 3× for 10 minutes with 20mM TRIS. The other 3 pieces were incubated for 20 minutes (T=20), 80 minutes (T=80) and 240 minutes (T=240) respectively, with 200μl 0.1U Nuclease *Bal 31* enzyme (New England Biolabs #M0213S) at 30°C, immediately followed by washing similar to T=0 to stop the digestion. After digestion, the TRF-protocol as described below was performed to measure the molecular size of TTAGG fragments.

### 2.5 Terminal Restriction Fragments protocol

We measured telomere length using similar procedures as in Boonekamp *et al.* (2014). Briefly: agarose plugs were pre-incubated for 1 hour with 1× Cutsmart buffer (New England Biolabs), followed by overnight incubation with 20U restriction enzymes HinfI and 20U RsaI (New England Biolabs) at 37°C. Telomeric repeats of sequence TTAGG were separated from restricted DNA with Pulsed Field Gel Electrophoresis (CHEF-DR II system by Bio-Rad) in a 0.8% agarose gel.

Electrophoresis settings were: 4.8V/cm, 1-25sec switch time, 22 hours at 14°C and for the *Bal 31* assay we used: 3.5V/cm, 0.5-7sec switch time, 24 hours at 14°C. Each gel also contained ^32^P-end-labeled size ladders (Molecular Weigh Marker XV, by Roche Diagnostics and 1Kb DNA ladder by New England Biolabs). After electrophoresis, the gels were fixed, dried and denatured. Denaturation was performed by incubating the dried gel with denaturation buffer (1.5M NaCl, 0.5M NaOH) 3 times for 30 minutes at room temperature, with gentle agitation, followed by neutralization buffer (0.5M Tris‐HCl pH 8.0, 1.5M NaCl) 2 times for 30 minutes at room temperature, with gentle agitation. Hybridization of telomeric repeats occurred overnight at 37°C with a ^32^P-end-labeled oligonucleotide (5’-CCTAA-3’)_4_ with gentle agitation in hybridization buffer (0.5M Na2HPO4, 7% SDS, 1mM EDTA). After washing away unbound oligonucleotides, the gel was exposed overnight to a phosphor screen (Perkin Elmer) and read at 600dpi with the Cyclone Storage Phosphor System by Perkin Elmer.

### 2.6 Statistical analyses

We analysed the effects of treatment on body mass and telomere length using mixed effects models with family ID as random effect. We corrected for gel differences in our telomere length analyses by including gel-ID as a fixed effect. We subsequently analysed temperature and age as fixed effects (ordinal factors) and the intercept therefore reflects the dependent variable at 23°C and at day 65. Mixed effects models were performed with the ASreml4 package in R using restricted maximum loglikelihood estimation. We used F-tests and log-likelihood ratio tests to determine the statistical significance of fixed and random effects, respectively.

We calculated the coefficient of variation of telomere length by dividing the square root of the residual variance (V_R_) of a model that only included gel-ID as a fixed effect with the mean telomere length 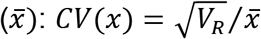. This is an appropriate estimate of biological variation because it corrects for methodologically induced variation and then scales the residual biological variation to the mean telomere length. We calculated the broad sense heritability (H^2^) of telomere length following Falconer & Mackay (1996) by dividing the between family variance (*V_f_*) estimated by the family ID random effect, with the total variation and then multiplying by two (since full siblings are on average related by 0.5): *H*^2^ = 2 ∙ *V*_*f*_/(*V*_*f*_ + *V*_*R*_).

## 3 RESULTS & DISCUSSION

*Bal 31* reduced telomere length in all 6 individuals tested for this purpose (Fig. 2). We found no evidence for the presence of large interstitial telomeric repeats as revealed by the absence of a banded pattern with single stranded DNA (Fig. 2). Our findings are in agreement with a recent FISH study in two other cricket species (Palacios-Gimenez *et al.* 2015). Together these findings support our assumption that our telomere length measurements reflect telomeric chromosome-ends.

**Figure 2.**
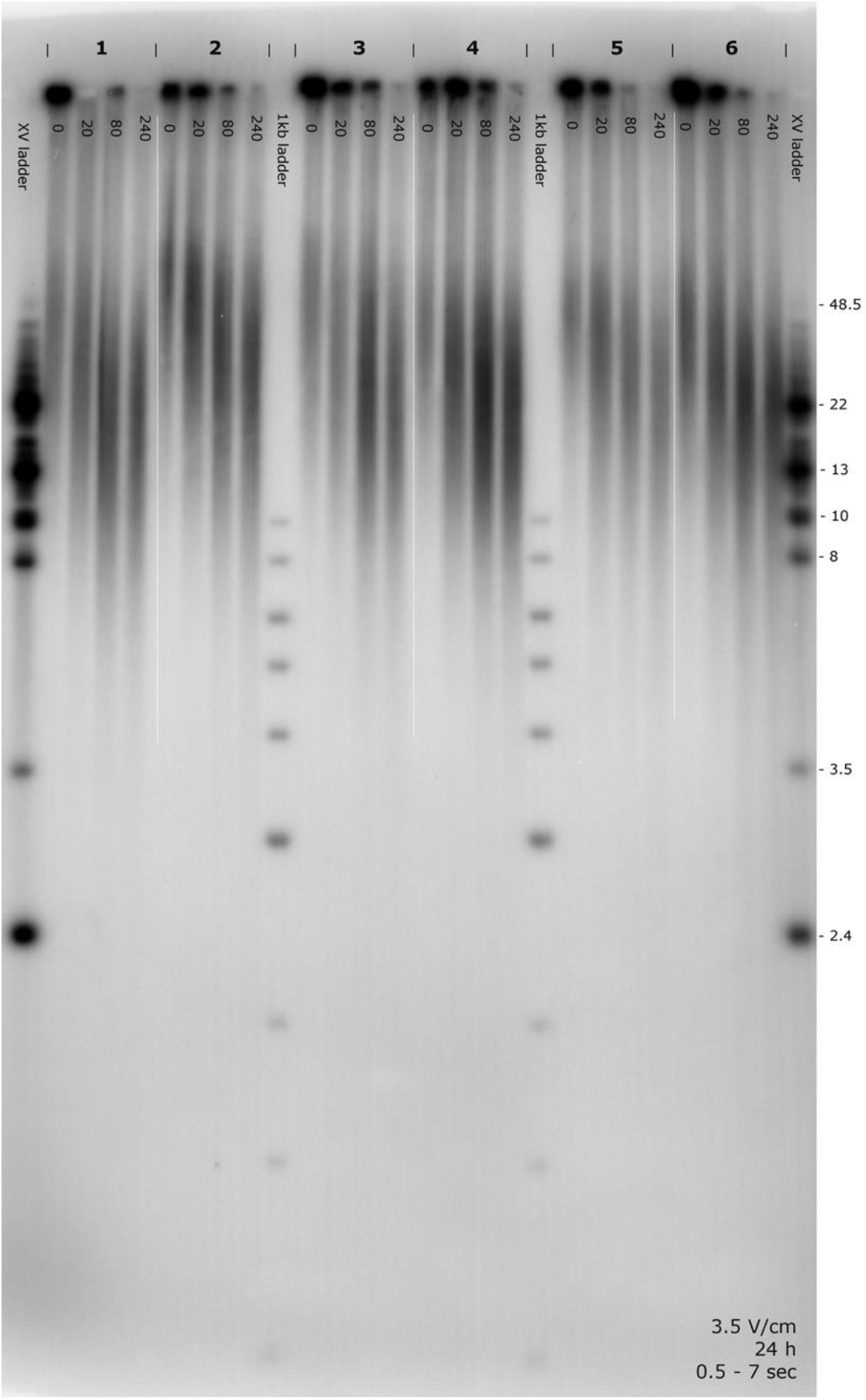
Pulsed field gel electrophoresis showing TTAGG_n_ telomere restriction fragments from the denatured chromosomes of six individual crickets (indicated by the numbers 1-6). XV molecular size ladders are shown on the outer lanes and shorter 1kb ladders are shown between crickets 2-3 and 4-5. The photo shows four aliquots of DNA sample per cricket and these aliquots were exposed to 0 (i.e. untreated), 20, 80, and 240 minutes of *Bal 31*. Each time series clearly shows that *Bal 31* decreases the molecular size distribution of TTAGG fragments. The absence of a banded distribution in the smears suggest that interstitial telomeric repeats were not present and that chromosome strands were intact. Taken together, these findings show that TTAGG_n_ fragments were located at the chromosome-ends.

Telomeres were on average 38 kb (mean= 38297) long, which is roughly 4 times longer that the telomere length of human newborns (Factor-Litvak *et al.* 2016), but in the same range as various other vertebrates e.g. (Gomes *et al.* 2011). The among-individual coefficient of variation was 10.3% and telomere length ranged from 27 – 50 kb.

Temperature treatment more than tripled the growth rate (Fig. 3A). On day 65, nymphs in the 23°C group were on average 0.12 ± 0.01 (s.e.) grams whilst nymphs in the 28°C group were 0.41 ± 0.01 grams (Table 2). Nymphs in the 23°C group required another 60 days to almost catch up at 0.35 ± 0.02 gram compared with the 28°C group at 65 days (Table 2). Nymphs from the intermediate age group of 85 days at 28°C were the heaviest (0.49 ± 0.01 grams).

**Table 2.**
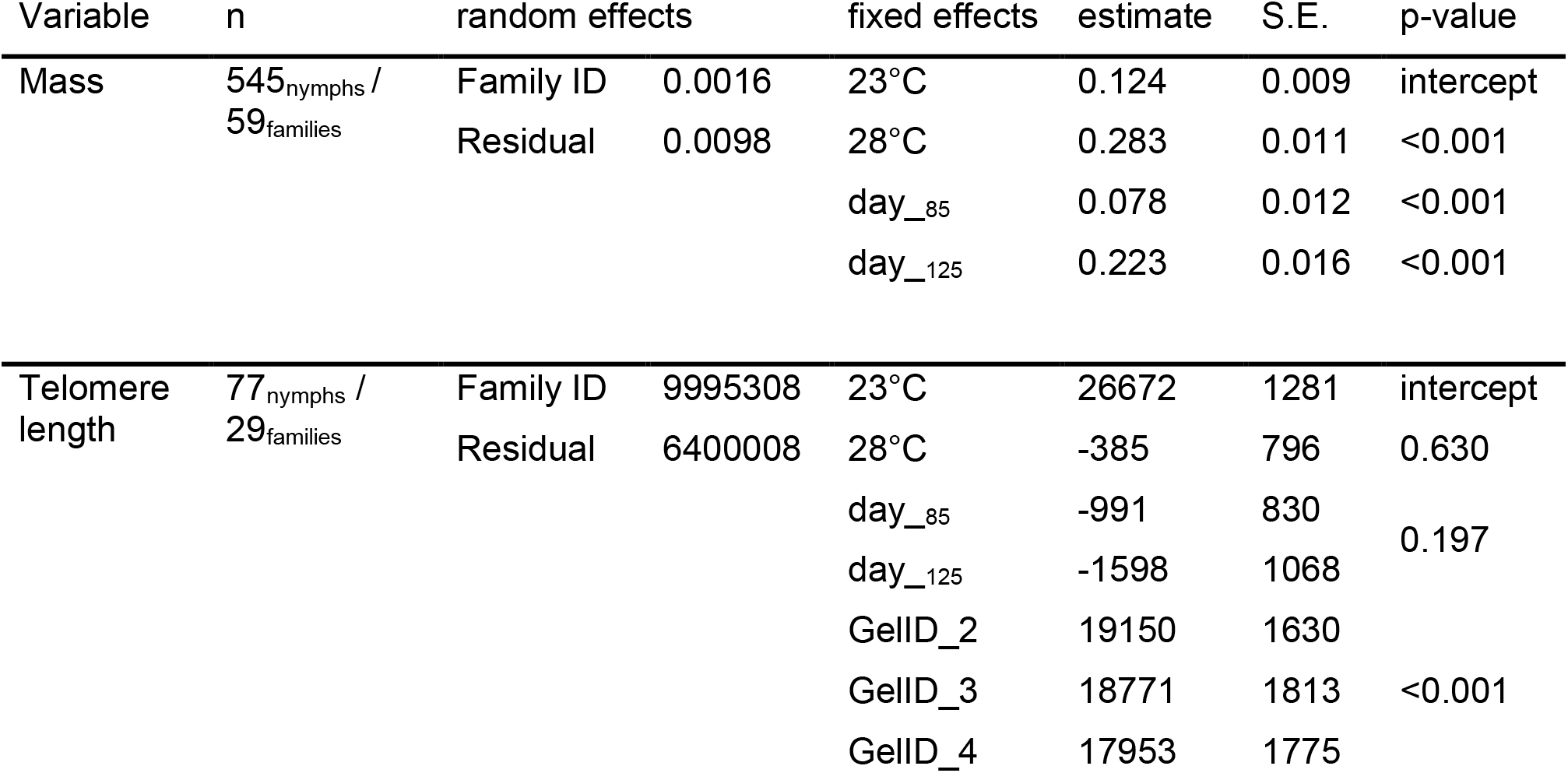
Body mass and telomere length in relation to temperature treatment and age. “n” reflects sample sizes at the individual and family levels. “random effects” shows the estimated family and residual variance components and “fixed effects” show the effects of temperature and time in treatment in days, which were included in the models as fixed factors. Note that the intercepts reflect body mass and telomere length at 23°C and 65 days old.

**Figure 3.**
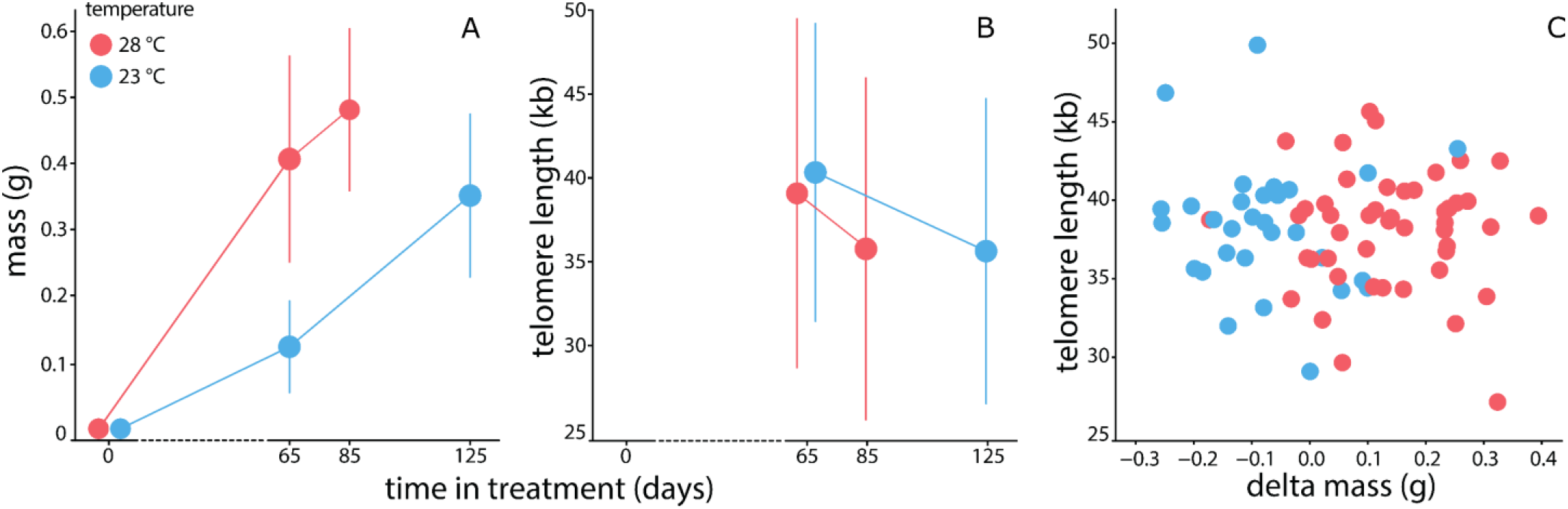
Body mass (A) and telomere length (B) of nymphs after 65-, 85-, and 125 days of temperature treatment. Nymphs started treatment when their body mass was virtually zero (included in the figure to graphically virtualise growth trajectories). Vertical bars are standard deviations. Delta mass (C) represents the individual deviation from the average mass of siblings.

Despite the strong effect on growth, temperature treatment did not significantly affect telomere length (Table 2; Fig. 3B). All four experimental groups showed indistinguishable mean telomere lengths, indicating that both temperature and chronological age had no significant effect on telomere length. Concomitantly, the overall correlation between body mass and telomere length was weak and non-significant (r = −0.074; p=0.52) and there was no evidence for an interaction between temperature and age, affecting telomere length. We also estimated the effects of average family mass and delta mass (i.e. the individual deviation from the family average) on telomere length, but neither of these mass components were significant (average mass = −16.5 ± 9.9 kb / g; delta mass = −2.5 ± 2.1 kb / g; Fig. 3C). The slope of age vs. telomere length analysed as continuous variable was 22bp/day (95% C.I. −5.67 – +50bp/day).

The random effect of family ID explained 61% of the total telomere length variation (Table2: p<0.001). This proportion was essentially unaltered by conditioning on the fixed effects of temperature and age (reflecting the fact that these effects explain negligible variance). The among family variance is remarkably high and indicative of high repeatability of telomere length among full siblings. This is an important observation, first because the high repeatability implies that our null findings with respect to temperature treatment and age were unlikely to be due to low measurement accuracy. Furthermore, the high repeatability on the level of full siblings potentially reflects high heritability. Following Falconer & Mackay (1996) we derived the broad sense heritability from the family mixed effects model by multiplying the full sibling repeatability by 2. Thus, the broad sense heritability of telomere length was 1.24 (95% C.I. 0.83 – 1.65). Graphical presentation of the data illustrates the strong correlation of telomere length among full siblings (Fig. 4). Applying the same calculations to body mass resulted in a broad sense heritability of 0.094 (−0.01 – 0.20) – or 0.14 (0.02 – 0.27), or 0.28 (0.11 – 0.45) conditional on including age or both age and temperature, respectively; this substantially lower heritability will arise, in part, from the strong effects of temperature and age that contribute to phenotypic variance in body mass.

**Figure 4.**
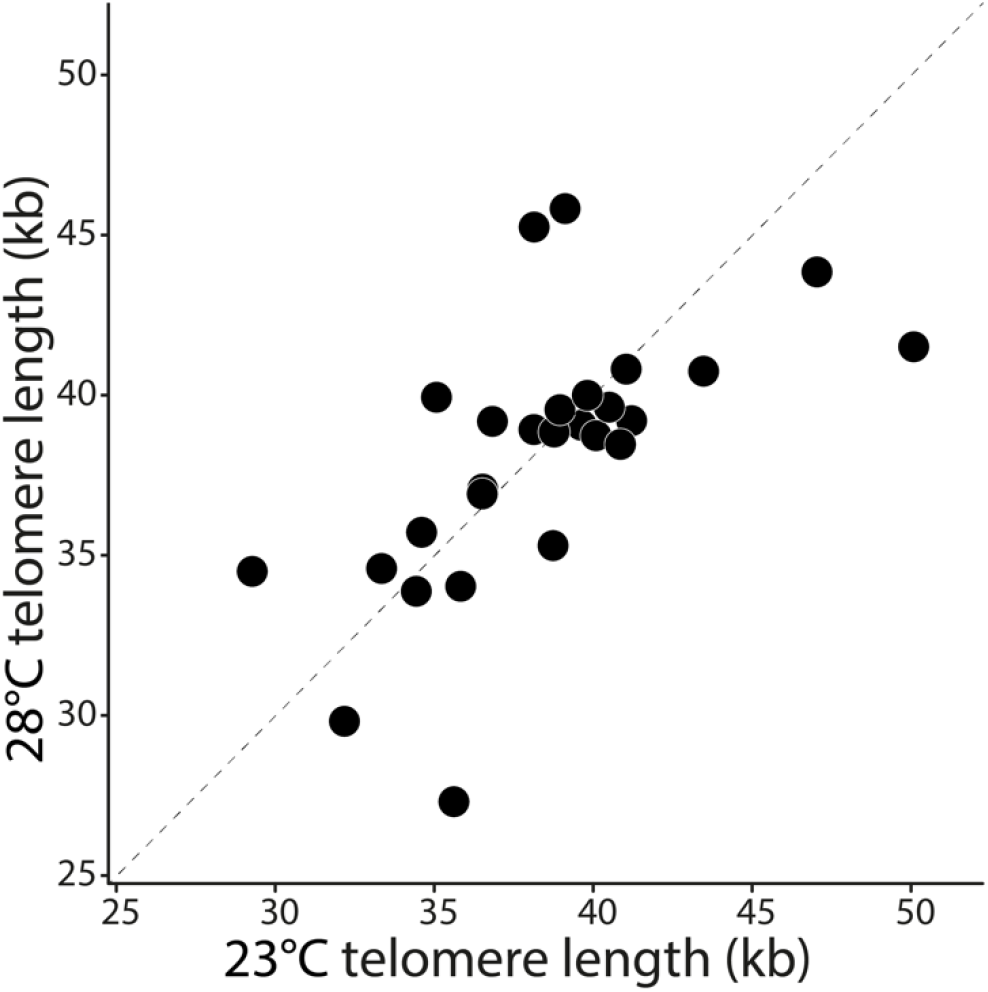
Telomere length of nymphs from the 23°C treatment group plotted against their siblings from the 28°C treatment group. The dashed line shows the line of equal values on the y– and x– axes. Note that deviation from the y=x line is expected even when the heritability is 100% since the genetic relatedness of full siblings is 0.5 on average.

Such a high heritability of telomere length intuitively suggests potential for rapid evolutionary change in response to selection. However, since heritability is a ratio, i.e. genetic variation relative to the total variation, this is not necessarily true. We therefore conducted additional post-hoc quantitative genetic analyses to directly estimate the additive genetic variance and evolvability (Houle 1992) of telomere length. We acknowledge that our study design is not optimal for this as the absence of any half-sibling structure means we cannot statistically separate additive genetic variance from any parental and/or dominance variance present. Noting that the results should thus be interpreted with caution, we fitted univariate animal models of telomere length and body mass to partition additive genetic and residual (environmental) variance components. Scaling to broad sense (or narrow sense if we assume an absence of parental and dominance variance) heritabilities corroborated our family-model estimates. For telomere length, heritability was ~100% with a standard error < 0.01 and the estimate was unaffected when conditioning the estimate on fixed effects of age and/or treatment. For body mass, we estimated a heritability (± s.e.) of 9.4±5.3%, increasing to 14.5±6.4% and 27.9±8.7% when age, or both age and temperature were included in the model as fixed effects.

We used the estimated (additive) genetic variance to calculate the coefficient of genetic variation: 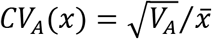, where *V_A_* is the (assumed additive) genetic variance of telomere length (*x*) that is estimated by the pedigree random effect. Following Houle (1992) and Garcia-Gonzalez *et al.* (2012), the evolvability is equal to the square of *CV_A_* and ranges from 0-100%. The telomere length coefficient of genetic variation was low (9.33%) resulting in a low evolvability of 0.87%. In comparison, the body mass coefficient of genetic variation was higher (19.06%) resulting in a higher evolvability of 3.64%, despite the lower broad sense heritability of 9.4%. Notably, the relatively low genetic variation on the mean standardised scale of telomere length is likely to limit its potential for evolutionary change under selection despite the high heritability.

Finally, we ran a bivariate animal model to determine the extent of genetic variation among genotypes in the response to temperature treatment (i.e. GxE). We did this to check the possibility of a GxE interaction resulting in the absence of an overall effect of temperature on telomere length (Fig. 3B). This model estimates the additive genetic variance within each temperature condition (*V_A23_* and *V_A28_*) and the genetic covariance (*cov_A_*) among them from which the genetic correlation can be derived: 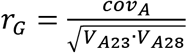. G × E interactions are observed either when *r_G_* is significantly below 1, and/or when the two additive genetic variances are significantly different. Age was included in both models as a fixed factor such that the observed genetic heterogeneity is over and above the effects of age. Gel ID was included as additional fixed factor for the telomere length model. We found that the genetic correlations (± s.e.) did not significantly deviate from 1 for body mass (*r_G_* = 0.62±0.27; *p* = 0.13) or telomere length (*r_g_* = ~1±0.01). However, for body mass *V_A23_* was significantly smaller than *V_A28_* (*V_A23_* = 0.0051 ± 0.0009; *V_A28_* = 0.0092 ± 0.0016; *p* = 0.044) supporting GxE. Graphical inspection of the data agreed with these analyses (Fig. 5).

**Figure 5.**
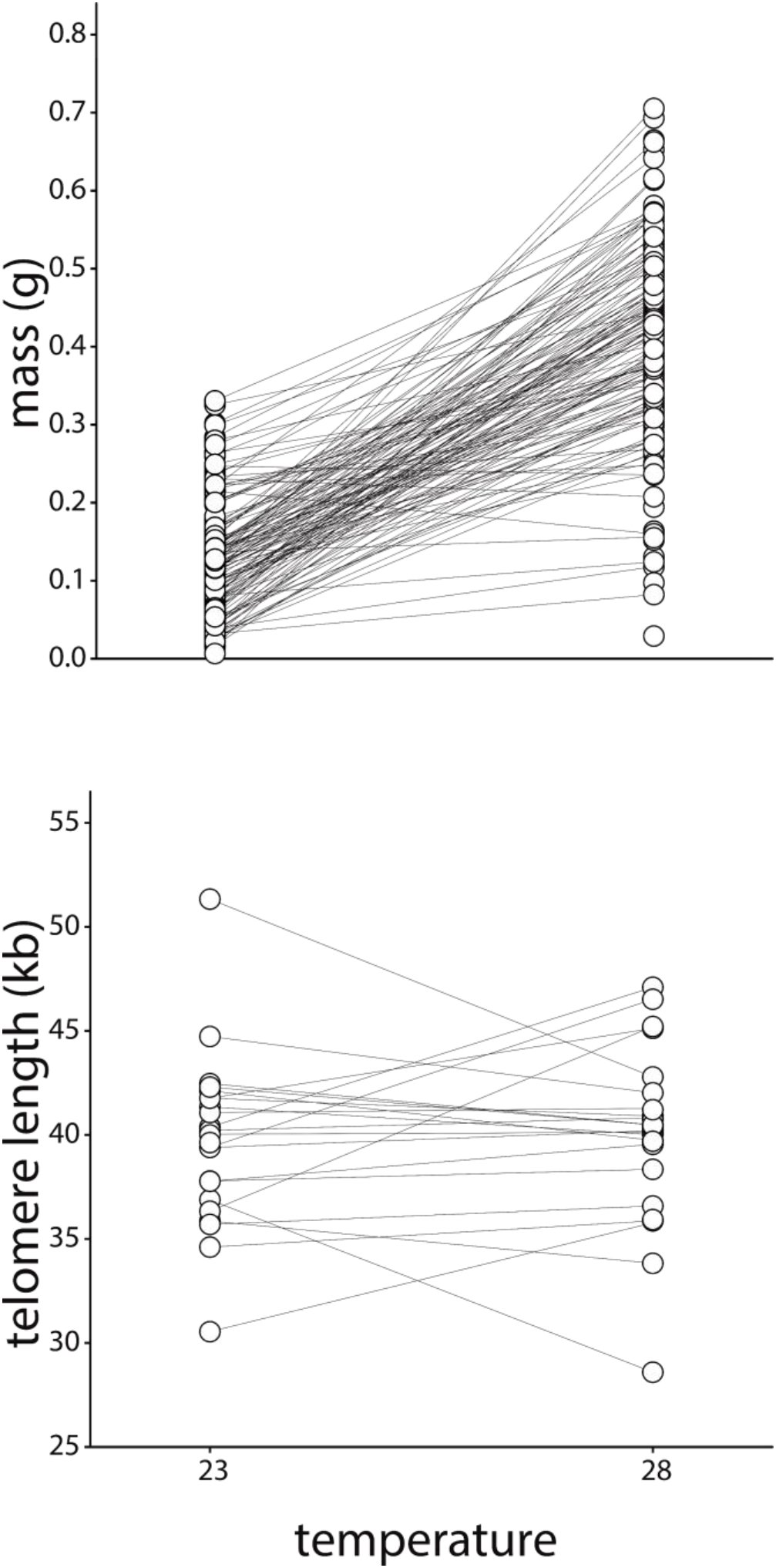
Body mass and telomere length in response to temperature treatment. Lines connect siblings. Note that for body mass, this figure only shows the 65 days after treatment subset because there were strong effects of age (in the statistical analyses we included all data and controlled for age). We included all telomere data in this figure since the effect of age on telomere length was not statistically significant.

In conclusion, our analyses do not support the hypothesis that in field crickets temperature induced growth acceleration is achieved at the cost of telomere length. However, in considering the implications of this finding (below) it is worth noting that the absolute mean telomere length difference between the two temperatures is 385 bp after 65 days of treatment (Table 2) which is substantial on an absolute scale when compared with previous studies that manipulated growth, e.g. Boonekamp *et al.* (2014). Likewise, the rate of telomere attrition is 22 base pairs per day, which approximately equals the annual telomere loss in adult humans. It is important to recognize that the size of the “effect” depends on what scaling is used and it remains unclear which scaling is the most informative with respect to fitness. For example, the effect size may be expressed as the mean difference between experimental groups relative to a common standard deviation, i.e. Cohen’s d (Nakagawa & Cuthill 2007). Given the high heritability it may also be useful to express the mean difference among treatments relative to the family residual variance thus expressing the effect of temperature on telomere length relative to variation within families. The first estimate results in a very low effect size of the temperature treatment (Cohen’s d = 0.04), whereas the latter approach results in a higher (albeit still small (Cohen 2013)) effect size of d = 0.19. It remains to be tested which scaling is more predictive of fitness and it could be that scaling to the family level variance could be more informative when this approach yields a better estimate of the rate of telomere attrition. A detailed study in the wild linking telomere length and the rate of telomere attrition to fitness is required to further our understanding of these variance scaling issues.

## 4 GENERAL DISCUSSION

Telomere dynamics have recently been proposed as a mechanism linking growth to fitness in vertebrates, e.g. Monaghan & Ozanne (2018). Metabolic rate and oxidative stress have been shown to increase with temperature in some insect species (Lalouette *et al.* 2011; Das *et al.* 2018) and, based on this, we expected growth acceleration by temperature to shorten telomeres, possibly through higher oxidative stress. However, we found that temperature-induced acceleration of growth had no discernible effects on telomere length near the end of the experimental growth period in field crickets. Our findings are in contrast to a comparable study showing that warmer temperatures accelerated pre-hatching growth and reduced telomere length in a bird species, the common tern *(Sterna hirundo)* (Vedder *et al.* 2018). A notable difference between the two study species is that crickets are ectotherms, and as such differ in their physiological response to environmental temperature from that of endotherms. At the same time, bird embryos are ectothermic during their early development and the role of ectothermy therefore remains speculative. It is interesting in this context that a study of another ectotherm, the Atlantic salmon (*Salmo salar*), also found no effect of temperature induced growth rate variation on telomere length (McLennan *et al.* 2018). These findings are unexpected because even if growth acceleration does not incur any fitness cost – as may be the case when growth is entirely temperature dependent – we would still expect the absolute number of cell divisions to affect telomere length. In contrast, we found that nymphs that were three to four times the size of their siblings had indistinguishable telomere length irrespective of the different age groups (Fig. 3). This raises the question of whether there could be mechanisms that actively maintain telomere length irrespective of age, size and growth rate, for example through the activity of telomerase (Collins & Mitchell 2002). From an evolutionary perspective, the answer to this question depends upon the level of phenotypic and genetic variation: In the case of low variation, robustness against environmental perturbations could indicate an adaptation because it suggests strong environmental canalisation of the trait (Waddington 1942; Stearns & Kawecki 1994; Boonekamp *et al.* 2017; Vedder *et al.* 2017; Boonekamp *et al.* 2018). In the opposite scenario, an almost complete absence of selection on the trait could have resulted in the accumulation of variation, but this variation would be neutral with respect to selection. Below, we discuss these possibilities.

A common explanation for high genetic variation in a trait is that selection is weak allowing mutations to accumulate. This is particularly easy to envisage in relation to telomere length, because telomere length represents the molecular size of non-coding DNA. Indeed, recent reviews of the literature have found that telomere length is highly heritable across vertebrates (Atema *et al.* 2015; Dugdale & Richardson 2018), including humans e.g. (Broer *et al.* 2013) and our observation fits this pattern. However, it remains unclear to what extent these empirical estimates of telomere length heritability reflect the amount of additive genetic variance, or alternatively, that the high heritability reflects low environmental variance (Hansen *et al.* 2011; Garcia-Gonzalez *et al.* 2012). Hence, it improves interpretability if estimates of genetic variance (and trait means) are provided alongside estimates of heritability (Morrissey *et al.* 2010). We used the coefficient of genetic variation because this fraction expresses the estimated additive genetic variance relative to the mean telomere length. This facilitates comparison of levels of genetic variation among traits and species with phenotypic values on different quantitative scales and is the recommended technique in evolutionary studies (Houle 1992). We recognize that the suitability of this approach for among-species comparisons hinges on whether the variance of telomere length scales to the mean across species, and to a lesser extent, whether such scaling of variance is comparable to that of other phenotypic traits. We are unaware of the existence of such quantitative comparative analysis. We found that the coefficient of genetic variation of body mass was about double that of telomere length, despite the much higher heritability of telomere length. The apparent low coefficient of genetic variation of telomere length is unlikely to reflect selective sampling because we sampled from 10 discrete, geographically isolated populations (Table 1).

If telomere length is a selectively neutral trait, we might expect to see greater levels of genetic variation than we observed for body mass, which we assume is not selectively neutral. Our finding of less genetic variation for telomere length suggests it is robust and maintained at its current level, consistent with adaptive developmental canalisation. The plausibility of this hypothesis depends on the assumptions (i) that telomere length causally affects fitness and (ii) that there are physiological mechanisms that maintain telomere length in the face of environmental perturbations:

i. Telomere length is predictive of survival in vertebrates, overall showing a 20% increased mortality risk per standard deviation of lower telomere length (Wilbourn *et al.* 2018). It is reasonable to hypothesize the same is true in insects, although this remains to be verified. We have demonstrated the existence of both actuarial and phenotypic senescence in the few weeks of the adult phase of the life cycle of our wild field crickets (Rodríguez-Muñoz *et al.* 2019b), and similar observations exist in a number of insects (Tatar *et al.* 1993; Bonduriansky & Brassil 2002; Hunt *et al.* 2004; Sherratt *et al.* 2011). Furthermore, *G. campestris* spend >70% of their life as growing nymphs, and from a theoretical perspective this extensive period of growth / cell divisions should not restrict the potential for nymphs to show substantial telomere attrition. The absence of an effect of manipulated growth on telomere attrition and the low level of genetic variation suggests that telomere length is actively maintained, consistent with the canalisation hypothesis. It is therefore of great interest to investigate potential mechanisms that could support the canalisation of telomere length.
ii. Recent studies have identified potential canalisation mechanisms for telomere length in vertebrates. The combined action of sister-chromatid exchange and co-localization of the DNA recombination proteins Rad50 and TRF1 has been found to cause telomere elongation during early embryogenesis (Liu *et al.* 2007). The observed genetic variation of telomere length therefore depends on the inheritance of such telomere elongation mechanisms during early embryogenesis, over and above the inherited initial telomere length of parental gametes. Furthermore, another study shows that telomeres that become too long during early development are trimmed back by telomeric zinc finger–associated protein expression (Li *et al.* 2017). These studies show that mechanisms are in place that elongate and shorten chromosome-ends during early development to create telomeres of a certain length. The high heritability of telomere length suggests that the joint action of these mechanisms is heritable, resulting in a canalised, developmentally robust pre-determined telomere length. At the same time epigenetic effects have also been observed showing that offspring telomere length decreases with paternal age (Bauch *et al.* 2019). The presence of such epigenetic effects implies that telomere length regulatory mechanisms may be limited in their capacity to fully compensate for the telomere length that was physically inherited from the paternal gametes. Furthermore, progressive telomere attrition may reduce the heritability of telomere length in old age, as has been found in humans (Bischoff *et al.* 2005) and this could reflect another constraint on the canalisation of telomere length caused by ageing, resulting in de-canalisation.

Functional explanations for a causal relationship between telomere dynamics and fitness are based on the premise that telomeres lose their protective function inducing cell cycle arrest when they erode to a critical limit (Steenstrup *et al.* 2017). We found that the rate of telomere attrition was 22 bp/day, which amounts to a daily loss rate of 0.06%. Although we do not know what the critical telomere length is in field crickets, this slope implies that telomeres of an average of 38kb would be fully eroded after 4.5 years (or 2 full years if we take the upper 95% confidence limit of the slope of age). This suggests that it is unlikely that there is a causal effect of telomere length limiting lifespan in this annual insect unless there are bouts of substantially accelerated telomere attrition during a later life stage. On the other hand, it should be noted that we used the average telomere length of a distribution of different telomere lengths within a sample/individual. The argument has been made that the chromosomes with the shortest telomeres cause senescence (Baird *et al.* 2003; Zou *et al.* 2004). The TRF methods we used allowed us to look at the shortest telomeres (i.e. the 5% percentile), which ranged from 2.7 kb to 24.5 kb (average±SD: 15±5.6 kb). This range suggests that for some individuals the shortest telomeres could easily be fully eroded within a year. Our findings therefore neither support nor refute a possible causal limit of telomere dynamics on the lifespan of this species.

Field crickets are an ancient species dating back to the Triassic (Song *et al.* 2015). Our observation that field crickets have their chromosome-ends capped by telomeres, just like vertebrate species, illustrates the strong evolutionary conservation of these chromosomal structures. It also raises the question of whether there are fundamental similarities or differences in telomere function between vertebrates and other groups. Recent syntheses propose that telomere length of vertebrates may be under stabilizing selection by functioning as an anti-cancer mechanism (Aviv *et al.* 2017; Young 2018). This hypothesis is primarily based on the observation that larger vertebrates with long periods of growth tend to have shorter telomeres than small vertebrates and the premise that this relationship coexists with a higher cancer risk in larger species, but see (Nunney *et al.* 2015). Shorter telomere length could be adaptive in large vertebrates when it selectively shuts down cell proliferation of unstable cells. Insects on the other hand have been suggested to be virtually free from cancer (Scharrer & Lochhead 1950) and this could be the basis for a fundamental difference between vertebrate and invertebrate groups in the link between growth and telomere dynamics. Consistent with this hypothesis we found that field crickets show surprisingly long telomeres of ~38 kb, much longer than many species of birds and mammals. However, detailed information on the occurrence of cancer is scant for wild insects and tumour growth has been documented in dung beetles (Scharrer & Lochhead 1950), which live much longer than most insects. Clearly, more studies are required on different insects and other invertebrates to determine systematic similarities and/or differences of telomere function among taxonomic groups.

## ACKNOWLEDGEMENTS

Caleb Peters, Emma May, Paola Leone and Zinnia Pennington collected most of the crickets used in this study. Anne Meurs aided with cricket husbandry, body mass measurements and snap-freezing of nymphs. This work was supported by the Natural Environment Research Council (NERC) standard grant NE/R000328/1 and the European Union’s Horizon 2020 research and innovation programme under the Marie Skłodowska-Curie grant agreement No 792215 (Boonekamp).

## AUTHOR CONTRIBUTIONS

This study was designed by JJB, SV, and TT. Adults were mated and eggs were collected and incubated by RRM and PH. JJB conducted the temperature manipulation experiment in Groningen. EZ and EM snap froze nymphs, measured body mass, and carried out the DNA extractions and telomere length analyses. *Bal 31* assays were done by JJB and EM. Statistical analyses were performed by JJB with help from AW, SV and TT. JJB wrote the first draft and all authors contributed to the final version.

## DATA AVAILABILITY STATEMENT

Data available from the Dryad Digital Repository: “Link XX to be added after final publication”

---

**Figure S1.**
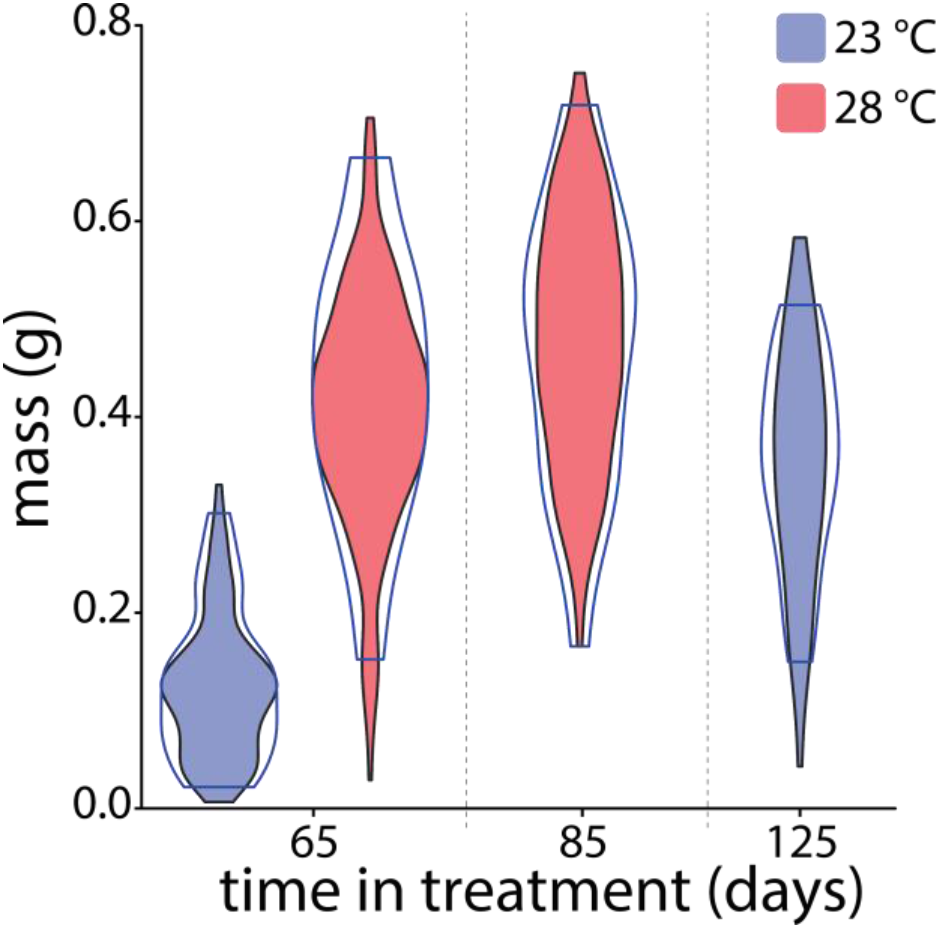
Violin plots of individual body masses measured before snap-freezing. Blue and red areas reflect masses at 23°C and 28°C respectively of all individuals. Contours reflect mass distributions of selected samples for telomere assay.

